# Elevated phosphate mediates extensive cellular toxicity: from abnormal proliferation to excessive cell death

**DOI:** 10.1101/2020.01.02.892638

**Authors:** Ping He, Olivia Mann-Collura, Jacob Fling, Naga Edara, Mohammed S. Razzaque

## Abstract

Inorganic phosphate (Pi) is an essential nutrient for human health. Due to our change in dietary pattern, dietary Pi overload engenders systematic phosphotoxicity, including excessive Pi related vascular calcification and chronic tissue injury. The molecular mechanisms of the seemingly distinct phenotypes remain elusive. In this study, we found that Pi directly mediates diverse cellular toxicity in a dose-dependent manner on a cell-based model. At moderately higher than physiological level, extracellular Pi promotes cell proliferation by activating AKT and extracellular signal-regulated kinase 1/2 (ERK1/2) cascades. By introducing additional Pi, we observed significant cell damage caused by the interwoven Pi related biological processes, including activation of mitogen-activated protein kinase (MAPK) signaling, endoplasmic reticulum (ER) stress, epithelial-mesenchymal transition (EMT) and apoptosis. Taken together, elevated extracellular Pi results in a broad spectrum of toxicity by rewiring complicated signaling networks that control cell growth, cell death, ER stress, and cell mobility.

## Introduction

Phosphate (PO_4_) is an essential nutrient for all living organisms. It maintains and modulates normal cell functions. *In vivo*, organic PO_4_ compounds are metabolized to inorganic phosphate (Pi) in the form of dihydrogen phosphate (H_2_PO_4_) and monohydrogen phosphate (HPO_4_). Pi serves as important substrate donor for important biological molecules, and plays key roles in a variety of biological processes, such as synthesis of DNA and RNA (as nucleic acid), storage and transfer of energy (as ATP), regulation of cell metabolism and cell signaling (as protein phosphorylation), and maintenance of cell membrane integrity (as phospholipids)(Michigami, Kawai et al., 2018). At systematic level in vertebrates, Pi is also critical for skeletal and dentin formation, growth and maintenance(Camalier, Yi et al., 2013).

Humans routinely consume phosphorus through food. Phosphorus is oxidized and transformed into PO_4_ after consumption within the body. Insufficient intake of dietary phosphorus, mostly due to malnutrition, can cause deficiency in skeletal mineralization, and development of rickets. In contrast, Pi overload accounts for the imbalance of phosphate metabolism leading to various human health problems, such as gingivitis(Goodson, Shi et al., 2019), heart disease(Cancela, Santos et al., 2012, Tonelli, Sacks et al., 2005), diabetes(Mancini, Affret et al., 2018), kidney disease(Kalantar-Zadeh, Gutekunst et al., 2010, Marks, Debnam et al., 2013) and cancer(Brown & Razzaque, 2018). Cellular studies have demonstrated extensive effects on both skeletal and extraskeletal tissues by elevated extracellular Pi. In skeletal cells, increase of Pi alters diverse cell behaviors, such as osteoblast(Beck, 2003) and osteoclast differentiation(Kanatani, Sugimoto et al., 2003), and vascular smooth muscle calcification(Giachelli, 2009). Extraskeletal, Pi has been identified as an essential nutrient for cell growth(Chang, Yu et al., 2006, Conrads, Yi et al., 2005) and proliferation(Roussanne, Lieberherr et al., 2001). In human lung cells, elevated Pi could accelerate cell growth by activating AKT and MAPK signaling(Chang et al., 2006), suggesting its potential roles in tumorigenesis. Studies in mice have suggested positive correlations between high serum Pi and cancer(Camalier, Young et al., 2010, Jin, Xu et al., 2009, Lee, Kim et al., 2015). In JB6 mice model, a high Pi diet resulted in cell transformation and skin tumorigenesis by stimulating N-Ras and downstream ERK1/2 phosphorylation(Jin et al., 2009). In K-ras^LA1^ mice, elevated Pi promoted lung cancer progression at early stage by stimulating cell proliferation and angiogenesis(Lee et al., 2015). On the other hand, evidence also showed Pi’s toxic effects of triggering cell death(Di Marco, Hausberg et al., 2008) and tissue damage(Nakatani, Sarraj et al., 2009, Yoshikawa, Yamamoto et al., 2018, Zhang, Yang et al., 2018). Abnormally high levels of Pi caused mitochondrial oxidative stress in endothelial cells, which further led to apoptosis(Di Marco et al., 2008). We previously showed elevated Pi (11.2±0.5 mg/dl) in *Fgf23*-null and *Fgf23/klotho* double-knockout mice compared to wild-type mice (7.7±0.3 mg/dl). The genetically induced high Pi in mice serum caused severe renal structure damage(Nakatani et al., 2009). Hyperphosphatemia was also found to be associated with skeletal muscle atrophy in mice mimicking chronic kidney disease (CKD)(Yoshikawa et al., 2018, Zhang et al., 2018). In humans, elevated dietary phosphorus consumption is associated with gingivitis by influencing cytokine levels(Goodson et al., 2019). Higher levels of serum phosphate are reported to be correlated with increased risk of heart failure and myocardial infarction(Tonelli et al., 2005). However, the molecular mechanism underlying the Pi induced tissue injury remains poorly understood.

In the present study, we comprehensively investigated Pi mediated cellular responses *in vitro*. First, we examined Pi’s effects on cell proliferation/viability by exposing cells to increase the amount of Pi. Next, we tested the alterations of various signaling transducers modulating AKT, MAPK, ER stress, EMT and apoptosis pathways in the context of high Pi. Finally, we applied inhibitors against these molecular events to the Pi treated cells to unravel how these biological processes interact and contribute to the broad spectrum of phosphotoxcity.

## Results

### 1. Elevated Pi increases proliferation and causes cell death in a dose-dependent manner

We reported excessive Pi could cause massive renal damage in mice that reduced overall survival (Nakatani et al., 2009). Literature also demonstrates the effects of Pi on cell proliferation in cells originated from kidney, such as HEK293(Yamazaki, Ozono et al., 2010). Here we also chose another immortalized cell line, HeLa, which is derived from cervical cancer and extensively described in literature. Both lines are widely used as homogenous experimental cell lines to ensure the consistency of cellular response and the reproducibility of data. Therefore, we used HEK293 cells as primary and HeLa cells as secondary cell models in this study. To study cell response to extracellular Pi, we firstly exposed HEK293 and HeLa cells to a wide range of Pi for 24hrs. The XTT assay analysis demonstrated that a moderate increase of extracellular Pi (up to 10mM) could promote cell viability/proliferation in HEK293 (Fig. 1A) and HeLa cells (Fig. 1B). The relative absorbance reached climax at 8mM of Pi by a 21% increase in HEK293 cells and a 24% increase in HeLa cells relative to the same concentration of sulfate (negative control) treated cells. When Pi concentration was above 16mM, it became detrimental (Fig. 1, A **and** B), and over 90% of the cells were dead at 64mM Pi. The data suggests that extracellular Pi may have a dose-dependent differential effect on cell proliferation and death.

**Fig. 1.**
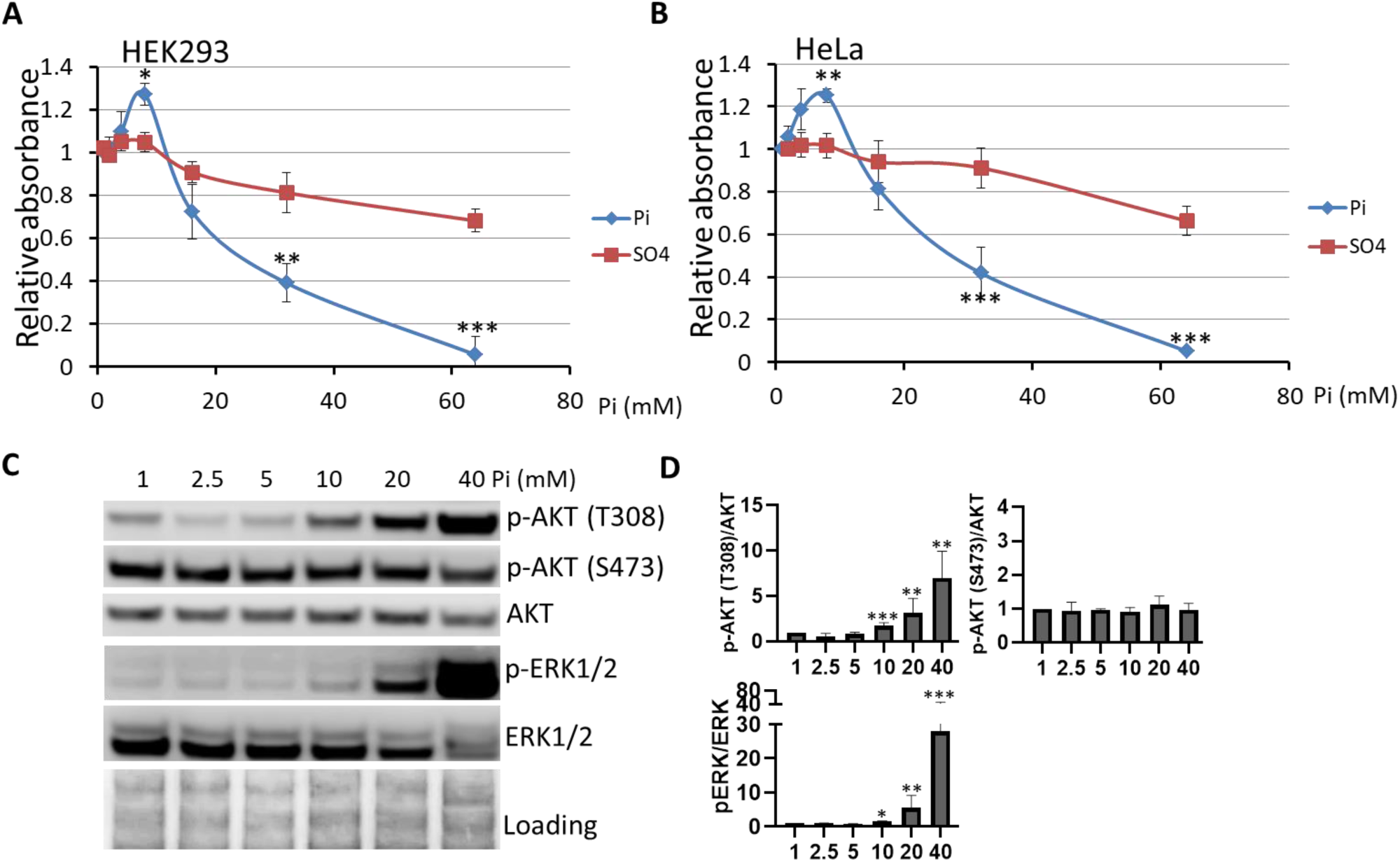
Phosphate has a dose-dependent differential effect on cell viability. HEK293 **(A)** and HeLa **(B)** cells were grown in DMEM culture medium (1mM Pi) with 10% FBS to 80-90% confluence followed by treatment with increased NaH_2_PO_4_ or Na_2_SO_4_ (negative control) for 24hrs. Cell viability was measured using XTT assay. The change in absorbance was measured on a plate reader 3hrs after the addition of XTT assay reagent. The data are plotted as the mean absorbance ± SEM (3 replicates/treatment) relative to no treatment. Unpaired *Student t*-test was used to compare means between NaH_2_PO_4_ and Na_2_SO_4_ treated groups. **(C) Moderately elevated extracellular Pi activates AKT and ERK signaling.** HEK293 cells were treated with an indicated amount of Pi for 24hrs. Total protein (15μg) extracted from each treatment was used for Western blot (WB) analysis with the anti-phospho AKT (T308), phospho AKT (S473), AKT, phospho ERK1/2 and ERK1/2 antibodies. The Ponceau S Stain of membrane was used as loading control. Bands quantification was done by densitometry image analysis **(D)** using Image Studio Lite software. The ratio of phosphorylated /Pan for each target is divided by that of the control (1mM Pi) to obtain the relative value for each treatment. Data represent means ± SEM from at least three independent experiments. Unpaired *Student t*-test was used to compare means between 1mM and higher Pi treated groups. \**P* < 0.05, \*\**P* < 0.01, and \*\*\**P* < 0.001.

High Pi exposure seems to control cell fate by balancing pro-survival/proliferation and pro-death mechanisms. We selected 1mM (normal culture condition), 10mM (pro-survival), 20mM (pro-survival/pro-death) and 40mM Pi (pro-death) Pi in the following immunofluorescence (IF) and Western blot (WB) analysis to investigate molecular mechanisms of Pi mediated cellular responses.

### 2. Moderately elevated extracellular Pi stimulates AKT and ERK signaling

To understand the molecular mechanisms of Pi related cell proliferation and cell death, we extracted the total proteins from cells treated with increasing amounts of Pi and analyzed target proteins level change by WB analysis. We found 10mM Pi treatment significantly augmented the phosphorylation of AKT at threonine 308 (T308), but did not alter the phosphorylation of AKT at serine 473 (S473) (Fig. 1, C **and** D). We also found slightly elevated phosphorylation of ERK1/2 (Fig.1, C **and** D). These data suggest a moderate increase of Pi could activate AKT and ERK signaling pathways.

### 3. Excessive Pi causes cell death, stimulates ER stress and EMT signaling

#### 3.1 High concentration of extracellular Pi triggers apoptosis

We observed high concentrations of Pi (over 16mM) could decrease cell viability in the XTT assay (Fig. 1, A **and** B). To understand the type of Pi induced cell death, we studied morphological changes in cell exposed to moderately high (10mM) and higher (40mM) levels of Pi under the microscope. IF microscopy showed that 40mM Pi treated cells, compared to non-treated and 10mM Pi treated groups, displayed clear cell shrinkage and fragmented nuclei in both HEK293 (Fig. 2A) and HeLa (Fig. 2F) cells. This represents the classic phenotypes of apoptosis. WB analysis further demonstrated the rise of cleaved Caspase-3/8 in the context of high Pi (20mM and 40mM) for 8hrs (fig. S1), 24hrs and 48hrs (Fig. 2, B, C, G **and** H). Necroptosis is another type of programmed cell death, which also involves the cleavage of Caspase-8(Gunther, Martini et al., 2011). We, therefore, tested the expression of two key necroptotic biomarkers, phosphorylated RIP and phosphorylated MLKL. However, we could detect neither of the proteins with high Pi exposure at times ranging from 8hrs to 48hrs (fig. S2, A **and** B).

**Fig. 2.**
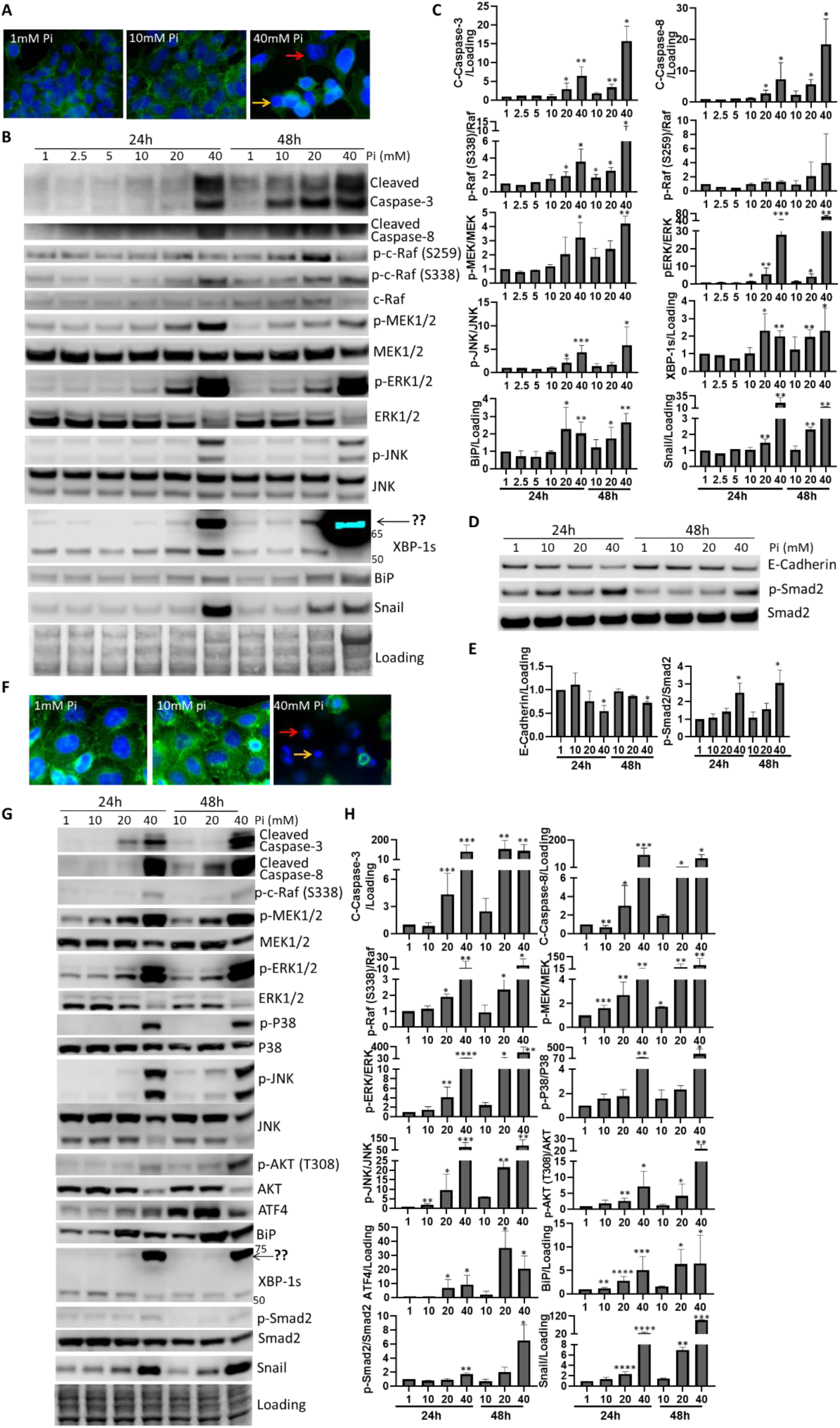
High concentration of Pi triggers apoptosis, activates MAPK, AKT, ER stress and TGF-β signaling. Immunofluorescence staining showing HEK293**(A)** and HeLa **(F)** morphology alterations when exposed to the indicated amount of Pi for 24hrs. Actin filaments were labeled with Alexa Fluor® 488 Phalloidin (green). Nuclei were counterstained with DAPI (blue). The red arrows indicate the fragmented nucleus, and the yellow arrows indicate the shrink cells. WB analysis (bands, **B, D, and G**, and quantitative analysis, **C, E and H**) shows high Pi induced apoptosis, MAPK, AKT, ER stress and EMT signaling in HEK293 **(B-E)** and HeLa cells **(G and H)**. Data represent means ± SEM from at least three independent experiments. Unpaired *Student t*-test was used to compare means between 1mM and higher concentrations of Pi treated groups. \**P* < 0.05, \*\**P* < 0.01, \*\*\**P* < 0.001, \*\*\*\**P* < 0.0001.

#### 3.2 Excessive Pi maintains the activation of AKT signaling and markedly enhances the MAPK signalling

While keeping concentration of extracellular Pi to 20mM and 40mM Pi, we observed a further increase of phosphorylation of AKT and ERK1/2, especially for the phosphorylation of ERK1/2 (Fig. 1, C **and** D). We then examined upstream signal transducers of ERK1/2 signaling, and detected extensive activation of the whole signaling pathway (inclined phosphorylation of c-Raf and MEK1/2) in the context of high Pi exposure (Fig. 2, B, C, G **and** H). Beyond ERK1/2 signaling, we tested other components of MAPK signaling molecules, such as Jun amino-terminal kinase (JNK) and p38. At higher concentrations of Pi, especially at 40mM Pi, we found a sharp increase of JNK and p38 phosphorylation (Fig. 2, B, C, G **and** H**, and** fig.S2C). These data indicate higher level (above 20mM) of Pi could further induce AKT and MAPK signaling.

#### 3.3 Elevated Pi leads to the activation of ER stress and EMT signaling

We speculated Pi overdosing may trigger homeostasis imbalance *in vivo* and consequently result in cell injury. ER stress is one of such key regulators of homeostasis(Walter & Ron, 2011). Hence, we examined the biomarkers of all three branches of ER stress pathways. We found increased levels of BiP, ATF4 and XBP-1s in high Pi (over 20mM Pi) treated cells compared to non-treated cells (Fig. 2, B, C, G **and** H). Interestingly, in 20mM and 40mM Pi treated HEK293 and HeLa cells, we detected a pronounced rise of modified XBP1s that shift its molecular weight (Mw) from 52kDa to 70kDa. This is possibly caused by post-translational modification (PTM). Unexpectedly, we also found TGF-β-induced EMT by showing a decline in E-Cadherin in HEK293 cells (Fig. 2, D **and** E) and an increase in phosphorylated Smad2 and Snail in both HEK293 (Fig. 2, B-E) and HeLa cells (Fig. 2, G **and** H). These results indicate elevated Pi could cause ER stress and induce EMT.

### 4. Interactions among the pathways in the context of high Pi

So far, our data demonstrate the direct link between excessive Pi and the following biological processes: activation of AKT, MAPK and TGF-β signaling, and enhancement of ER stress, EMT and cell death. We wanted to further study how extracellular Pi orchestrates these elements to contribute to the pro-survival and pro-death effects. To address this fundamental question, we applied inhibitors of AKT, MEK1/2, JNK and ER stress to the cultured cells exposed to increase amount of Pi.

#### 4.1 AKT inhibitors diminish cell proliferation/survival by repressing mTOR, and activating ERK1/2 and ER stress in high Pi settings

MK-2206 is a highly selective AKT inhibitor with high potency(Hirai, Sootome et al., 2010). Mammalian target of rapamycin (mTOR) Complex 2 (mTORC2) can phosphorylate and activate AKT(Saxton & Sabatini, 2017). Torin 1, a potent inhibitor of mTORC1/2(Thoreen, Kang et al., 2009), can therefore indirectly block AKT activity. We found these two AKT inhibitors could successfully suppress AKT phosphorylation (Fig. 3, A-D, and fig. S3), and they further abrogated high Pi upregulated mTOR signaling downstream of AKT by markedly attenuating the phosphorylation of P70S6K and RPS6 (Fig. 3, A **and** B).

**Fig. 3.**
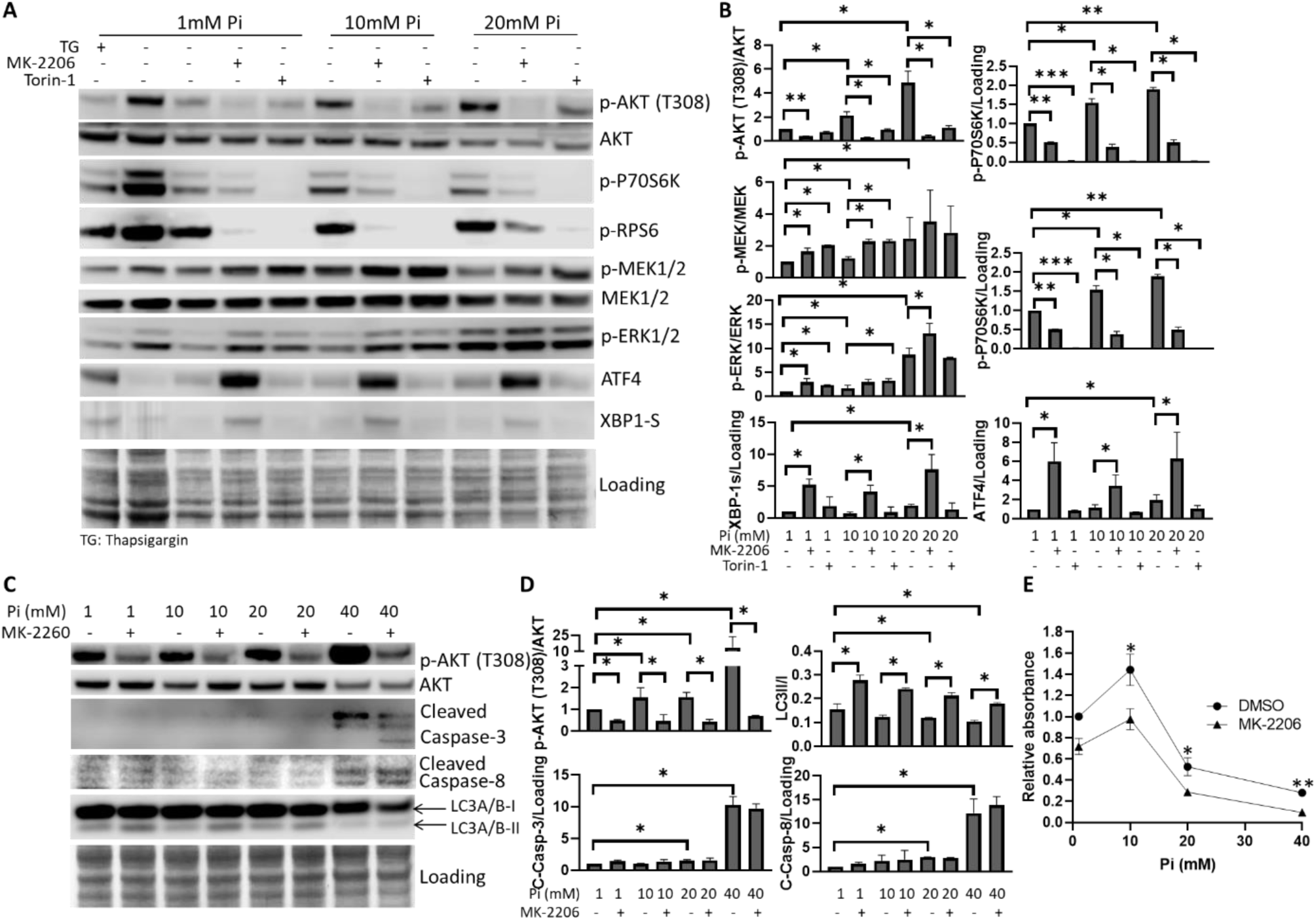
AKT inhibitors aggravates Pi toxicity by inactivating AKT-mTOR signaling, activating ERK signaling and ER stress in HEK293 cells. WB **(A)** and quantitation **(B)** analysis shows 5µM MK-2206 and 0.5µM Torin-1 suppressed AKT-mTOR signaling, activated ERK signaling and ER stress in the context of 10mM and 20mM Pi. Thapsigargin (TG) is ER stress enhancer, used as positive control (at concentration of 0.5µM) for ER stress. WB **(C)** and quantitation **(D)** analysis shows MK-2206 alleviated Pi suppressed autophagy (reduced ratio of LC-II/LC-I), but had no effect on apoptosis. **(E)** XTT assay demonstrates MK-2206 abolished 10mM Pi promoted cell proliferation and exacerbated higher Pi (20mM and 40mM) resulted cell loss. Data represent means ± SEM from at least two independent experiments. Unpaired *Student t*-test was used to compare means between 1mM and higher concentrations of Pi treated groups. Paired *t*-test was used to compare means between non-treated and MK-2206 treated groups at indicated concentrations of Pi. \**P* < 0.05, \*\**P* < 0.01, \*\*\**P* < 0.001.

Conversion of LC3-I to LC3II is a commonly used marker to reflect autophagic flux(KlionskyAbdelmohsen et al., 2016). High Pi at 20mM and 40mM concentrations blocked autophagy presented as the reduction of LC3II/I ratio by 23% and 35% respectively, in contrast to 1mM Pi treated cells (Fig. 3, C **and** D) suggesting excess Pi mitigated autophagy. To facilitate cell growth, AKT-mTOR signaling slows down the protein turnover by suppressing autophagy(Saxton & Sabatini, 2017). MK-2206, acting as an autophagy enhancer here, restored autophagy as seen in elevated in LC3II/I ratios in various concentrations of Pi treated cells (Fig. 3, C **and** D**, and** fig. S3). This indicates Pi repressed autophagy is dependent on the overstimulated AKT-mTOR cascades.

In addition, the XTT assay demonstrated that MK-2206 significantly attenuated cell growth by 28% at 1mM Pi, abolished Pi promoted cell proliferation at 10mM Pi and exacerbated Pi-elicited cell loss by 47% at 20mM and 67% at 40mM (Fig. 3E). However, we did not observe significant change in Cleaved Caspase-3/8 (Fig. 3, C **and** D) suggesting high Pi activated AKT signaling does not affect apoptosis. More interestingly, AKT inhibitors, especially MK-2206, also distinctively enhanced the phosphorylation of MEK1/2 and ERK1/2, and elevated the expression of ATF4 and XBP1s (Fig. 3, A **and** B), indicating AKT’s inhibitory impact on ERK1/2 signaling and ER stress. However, MK-2206 did not seem to perturb TGF-β induced EMT in HEK293 and HeLa cells (fig. S4).

**Fig. 4.**
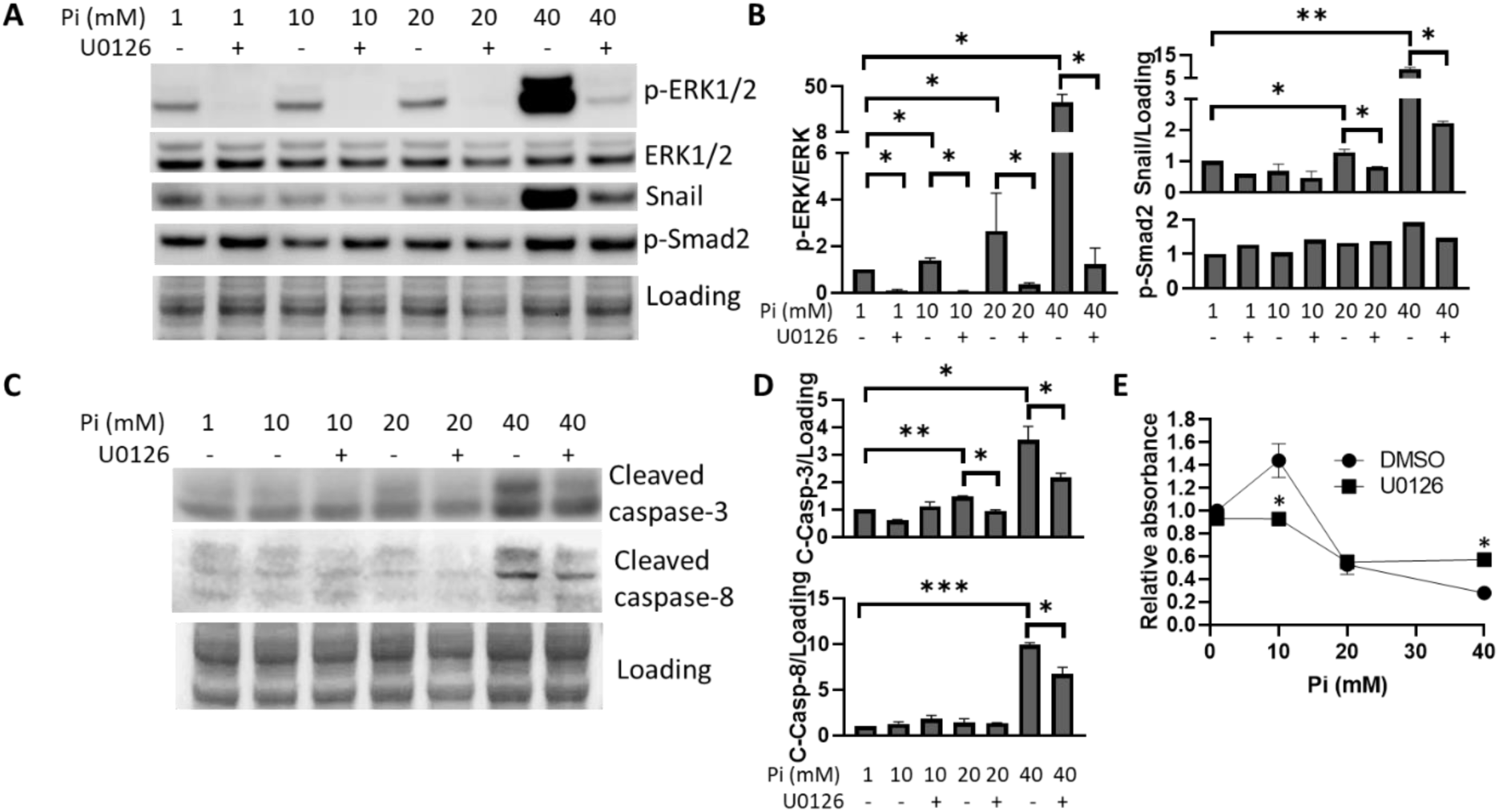
MEK inhibitor represses cell death and Snail up-regulation due to high Pi treatment in HEK293 cells. WB analysis shows MEK inhibitor, U-0126, at concentration of 10µM, inhibited ERK signaling, Snail up-regulation **(A and B)** and apoptosis **(C and D)**. **(E)** XTT assay shows10µM U-0126 prevented 40mM Pi resulted in cell death. Data represent means ± SEM from at least two independent experiments. Unpaired *Student t*-test was used to compare means between 1mM and higher concentrations of Pi treated groups. Paired *t*-test was used to compare means between non-treated and U-0126 treated groups at indicated concentrations of Pi. \**P* < 0.05, \*\**P* < 0.01, \*\*\**P* < 0.001.

#### 4.2 MEK1/2 inhibitor represses high Pi exposure induced cell death and EMT signaling

U0126 is a highly selective inhibitor of MEK1 and MEK2(Favata, Horiuchi et al., 1998). It abolished 40mM Pi boosted phosphorylation of ERK1/2 (Fig. 4, A **and** B), and in turn partially prevented Pi-induced cell death (Fig. 4, E) by reducing Cleaved Caspase 3/8 (Fig. 4, C **and** D). This suggests that Pi associated cell death is partially ascribed to hyper-stimulated ERK1/2 signaling promoted apoptosis. U0126 treatment also in part blunted excessive Pi induced upregulation of Snail, but it did not alter the phosphorylation of Smad2 (Fig. 4, A **and** B**, and** fig. S5) indicating that high Pi stimulated ERK1/2 signaling increases the expression of Snail via a Smad2 independent mechanism.

**Fig. 5.**
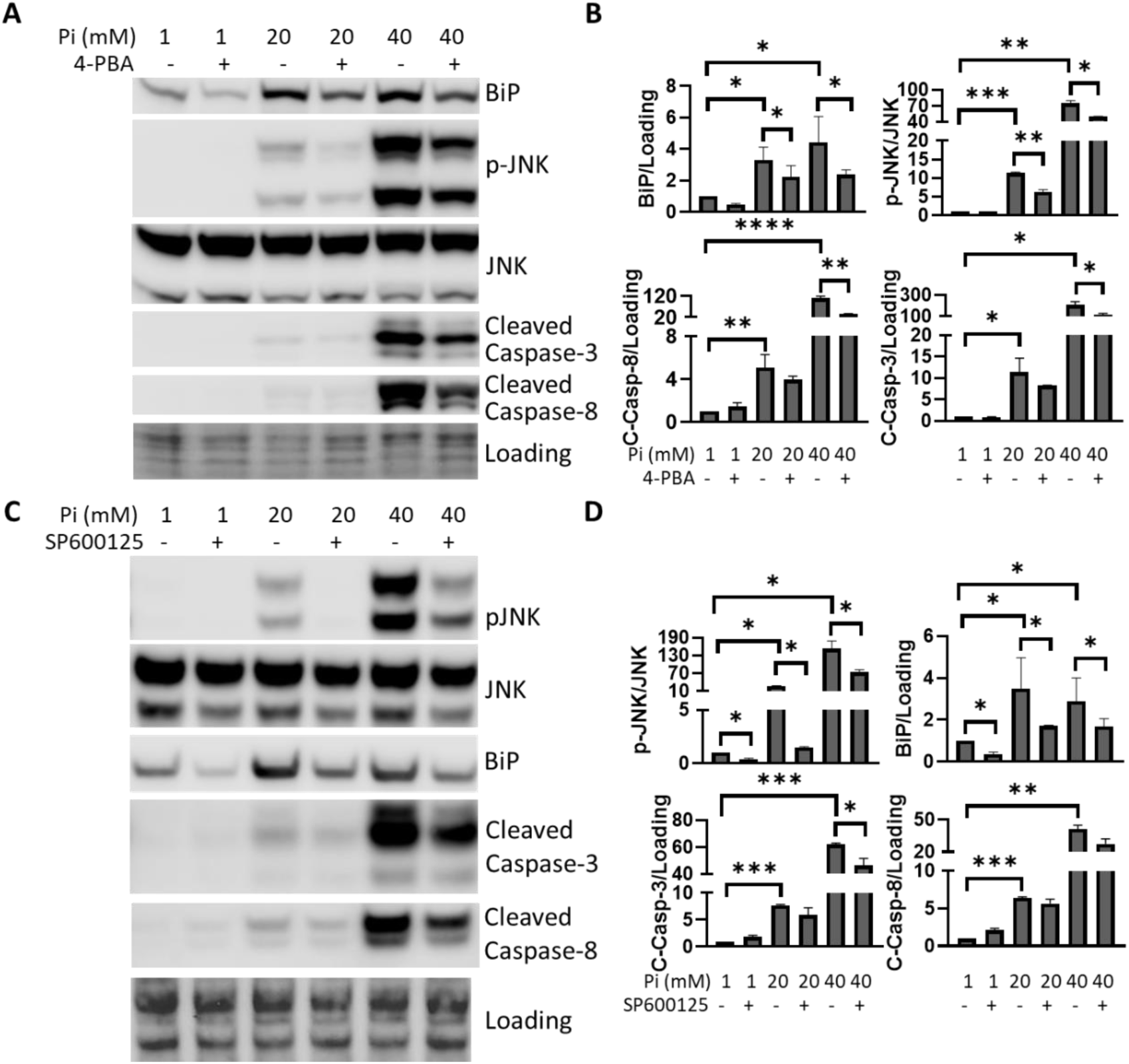
ER stress and JNK inhibitors reduces high Pi caused apoptosis in HeLa cells. WB **(A)** and quantitation **(B)** analysis shows ER stress inhibitor, 4-PBA, at concentration of 1µM, blunted high Pi mediated ER stress and Caspase-3/8 cleavage. WB **(C)** and quantitation **(D)** analysis shows JNK inhibitor, SP600125, at concentration of 50µM, abrogated high Pi enhanced JNK phosphorylation, BiP expression and cleavage of Caspase-3/8. Data represent means ± SEM from at least two independent experiments. Unpaired *Student t*-test was used to compare means between 1mM and higher concentrations of Pi treated groups. A paired *t*-test was used to compare means between non-treated and 4-PBA or SP600125 treated groups at indicated concentrations of Pi. \**P* < 0.05, \*\**P* < 0.01, \*\*\**P* < 0.001, \*\*\*\**P* < 0.0001.

#### 4.3 ER stress and JNK inhibitors reduce high Pi caused apoptosis

4-Phenylbutyric acid (4-PBA), acting as a chemical chaperone, can reduce the load of unfolded proteins retained in the ER during ER stress(Qi, Hosoi et al., 2004). 4-PBA significantly attenuated the level of high Pi upregulated ER stress sensors, such as BiP and phosphorylated JNK (Fig. 5, A **and** B). The ER stress inhibitor also repressed high Pi mediated cleavage of Caspase-3 suggesting its role in anti-apoptosis. In parallel, a JNK inhibitor, SP600125, could not only impede excessive Pi mediated activation of JNK signaling, but distinctively diminished BiP level as well (Fig. 5, C **and** D). We also observed SP600125 caused a decline of Cleaved-Caspase 3/8 indicating its role in preventing apoptosis. The data reveal that excessive Pi induced persistent ER stress and JNK signaling may contribute to cell apoptosis.

## Discussion

Pi has been identified as an essential nutrient for cell growth(Chang et al., 2006, Conrads et al., 2005) and proliferation(Roussanne et al., 2001). Its potential roles in tumor development have been implicated in in cancers of breast(Lin, McKinnon et al., 2015), lung(Jin et al., 2009, Lee et al., 2015), skin(Camalier et al., 2010) and kidney(Brown & Razzaque, 2018). Paradoxically, lines of evidence also demonstrate Pi overload has systematically detrimental effects such as cardiovascular disorders(Tonelli et al., 2005) and renal dysfunction(Marks et al., 2013). We found an increased level of Pi may have dose-dependent effects on cells. Moderately augmented extracellular Pi (1mM-10mM) may promote cell growth and proliferation, while further increasing the concentration of Pi (above 20mM) triggered cell death (Fig. 1, A **and** B).

The serine/threonine kinase AKT/PKB is one of the most critical signaling nodes within mammalian cells(Manning & Toker, 2017). Growth factors could activate it via PI3K and PDK1 by phosphorylating AKT at T308, which facilitates its phosphorylation by mTORC2 at serine 473 (S473). Activated AKT then phosphorylates reams of downstream signal transducers that govern cell survival, cell cycle progression, mobility and metabolism(Fruman, Chiu et al., 2017). Chang et al. reported Pi stimulated AKT phosphorylation on-site T308 specifically, which in turn triggered phosphorylation of RAF, MEK and ERK, indicating Pi’s involvement in cell proliferation(Chang et al., 2006). Consistently, we observed similar phenomena at pro-survival dosage of Pi (Fig. 1, C **and D,** Fig. 2, B **and** C), and Pi promoted cell proliferation was blunted by AKT inhibitor (Fig. 3E). mTORC1, activated downstream of PI3K-AKT signaling, is one of the key regulators of metabolism and growth(Dibble & Cantley, 2015). We verified high Pi stimulated mTOR signaling (Fig. 3, A **and** B). AKT inhibitors treatment data further reveal the pro-survival role of AKT signaling during the process of high phosphate-induced cytotoxicity. The Pi stimulated AKT signaling could keep cell viable/proliferating by activating its downstream mTOR signaling and blocking autophagy, ERK1/2 signaling and ER stress (Fig. 3 **and** Fig. 6).

**Fig. 6.**
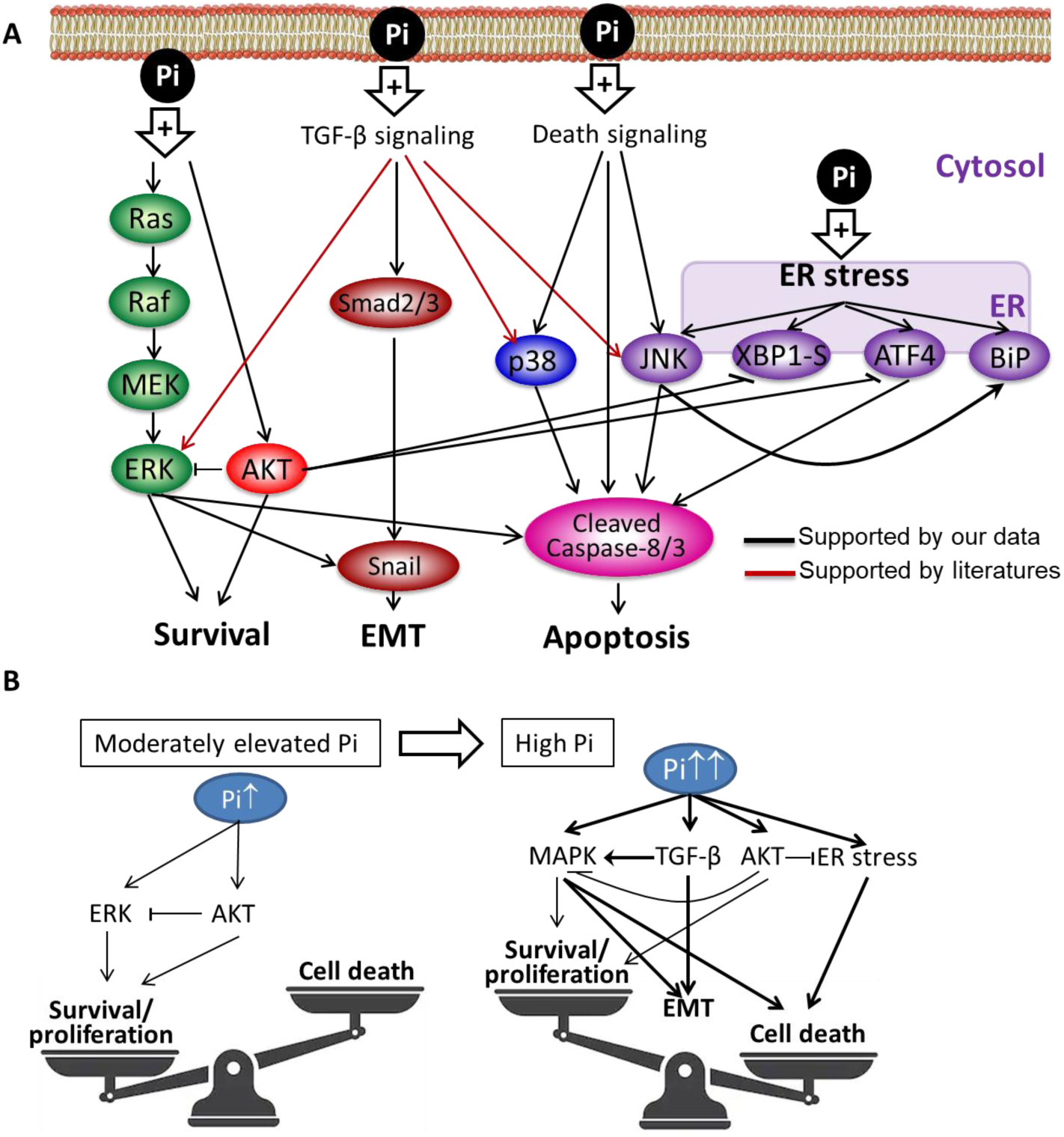
Schematic diagram of cellular responses to elevated Pi *in vitro.* **(A)** Pi rewired cell signaling networks. **(B)** Pi’s dose-dependent differential effects on cellular responses.

MAPK families, composed of ERKs, JNKs and p38/SAPKs (stress-activated protein kinases)(Zhang & Liu, 2002), play essential roles in regulating cell proliferation, differentiation and cell death in mammals(Morrison, 2012). A typical RAS–ERK signaling cascade involves subsequent activation of RAS, RAF, MEK1/2 and ERK1/2, and predominantly promotes cell growth, differentiation and cell cycle progression (Fig. 6A)(Lavoie & Therrien, 2015). Studies show that elevated extracellular Pi could activate the ERK signaling pathway in various mammalian cell types, but there seems to be a cellular specificity(Michigami et al., 2018). Beck and colleagues reported that 10mM extracellular Pi induced *Opn* gene expression in osteoblastic MC3T3-E1 cells through the stimulation of ERK1/2 pathway but not that of p38 or JNK signaling pathways(Beck & Knecht, 2003). Studies also showed that Pi stimulated RAF/MEK/ERK pathway favored cell proliferation by upregulating the levels of matrix Gla protein(Julien, Magne et al., 2007), Cyclin D1 and alkaline phosphatase(Kimata, Michigami et al., 2010) in chondrogenic cells, dentin matrix protein 1 (Dmp1), fibroblast growth factor 2 (Fgf2), Fgf receptor 1 (Fgfr1) and early growth response 1 (Egr1) in osteoblastic cells(Nishino, Yamazaki et al., 2017), and *EGR1* in HEK293 cells(Yamazaki et al., 2010). Conversely, several studies reported ERK signaling associated cell death via intrinsic and extrinsic apoptotic pathways(Cagnol & Chambard, 2010). We observed both anti- and pro-apoptotic functions of ERK signaling. Moderate increase of extracellular Pi to 10mM slightly enhanced ERK1/2 signaling (Fig. 1, C **and** D). A MEK1/2 inhibitor, U0126, mitigated Pi promoted cell proliferation (Fig. 4E), suggesting ERK pathway’s contribution to cell survival. The introduction of additional Pi to 20mM and 40mM markedly activated ERK and subsequently led to apoptosis by activating Caspase-3/8 (Fig. 2, B **and** G). U0126 blocked apoptosis triggered by high Pi stimulated ERK signaling (Fig. 4), indicating the direct link between high Pi upregulated ERK signaling and cell death. Unlike ERKs pathways, environmental stresses (such as ER stress and oxidative stress) and inflammation activated JNK and p38 are mainly involved in apoptosis and inflammatory response(Morrison, 2012). Indeed, we observed a dramatic induction of JNK and p38 phosphorylation at 40mM Pi (Fig. 2, B, C, G **and** H, fig. S2C). JNK inhibitor could partly prevent the cleavage of Caspase-3/8, indicating that activated JNK also contributes to excessive Pi caused apoptosis. Thus, Pi activated ERK signaling may serve as a “double-edged sword”. At moderately high level of Pi, it keeps cell growth and survival together with Pi upregulated AKT signaling. The introduction of additional Pi further stimulated ERK, which gave rise to cell death by functioning synergistically with other members (JNK and p38) of MAPK.

The vast majority of secreted and plasma membrane proteins enter the ER lumen first to fold and assemble(Hetz, Martinon et al., 2011). Various physiological and pathological conditions disrupt protein folding and result in the accumulation of unfolded or misfolded proteins. This cellular condition is termed ER stress, which triggers a series of adaptive reactions by activating intracellular signaling pathways termed the unfolded protein response (UPR)(Walter & Ron, 2011). Three principal signaling pathways of the UPR, mediated by PKR-like ER kinase (PERK), transcription factor 6 (ATF6), and IRE-1, function in parallel to increase ER protein-folding capacity, decrease protein load and restore ER homeostasis. If these mechanisms of adaptation fail to reestablish ER homeostasis, cells would die by the mechanism of apoptosis(Iurlaro & Munoz-Pinedo, 2016). We found high Pi associated upregulation of ER stress markers (Fig. 2, B, C, G **and** H) falling into three major branches of UPR (BiP from ATF6 branch, ATF4 from PERK branch, and phosphor-JNK, XBP1-s from IRE-1 branch), confirming ER stress as one of the consequences of high Pi exposure. Interestingly, WB analysis showed an extra band of XBP1s in high Pi treated cells suggesting a Pi associated PTM of XBP1s. Jiang, et al. reported a similar PTM of XBP1s (15-20kDa increase in Mw), and demonstrated it as a SUMOylated XBP1s. The investigators further showed the accumulating XBP1 SUMOylation increased ER stress-induced apoptosis(Jiang, Fan et al., 2012). In context of high Pi, elevated SUMOylated XBP1s may be one of the contributing factors to cell death. However, further studies are needed to figure out the nature of Pi associated XBP1s modification. Finally, ER stress inhibitors could partially block apoptosis (Fig. 5, A **and** B) indicating ER stress participates in high Pi-induced cell damage.

High Pi environment could also potentiate cancer metastasis(Brown & Razzaque, 2018). CKD-induced hyperphosphatemia is correlated with depressed Klotho expression(Kuro, 2011). Alpha-klotho maintains phosphate homeostasis and serves as a tumor suppressor in many types of cancers(Wolf, Levanon-Cohen et al., 2008). Klotho can suppress metastasis by inhibiting transforming growth factor-beta (TGF-β)(Doi, Zou et al., 2011). However, there is no concrete evidence showing a direct linkage between Pi and metastasis. We focused on TGF-β signaling induced EMT, which promotes cancer metastasis by reducing intercellular adhesion (decrease in E-Cadherin) and increasing motility (increased Smad2/3 phosphorylation and Snail)(Hanahan & Weinberg, 2011). *In vitro,* high environmental Pi reduced expression of E-Cadherin and increased level of phosphor-Smad2 and Snail (Fig. 2, D, E, G **and** H) indicating high Pi induced TGF-β signaling and EMT. More *in vivo* studies are desired to confirm Pi related TGF-β signaling and EMT in tumor progression. There is also extensive cross-talk between MAPK and TGF-β signaling, as reviewed by Gui, et al (Gui, Sun et al., 2012). For instance, TGF-β can activate the whole MAPK family (ERK, p38 and JNK) in various cell types via Smad dependent and -independent transcriptional mechanisms. Hence, Pi activated MAPK signaling may be in part ascribed to Pi upregulated TGF-β signaling. Meanwhile, ERK signaling can enhance EMT via transcriptional upregulation of Snail. Indeed, we found ERK signaling inhibitor could blunt high Pi-induced Snail (Fig. 4, A **and** B), confirming the ERK’s positive effects on EMT in the context of high Pi (Fig. 6).

To further study the Pi regulated signaling networks, we are currently expanding our research to various other primary and cancer cells derived from kidney, skin, and lung. We have provided clear evidence of high Pi-induced cell death, and shown the signaling cross-talks that are involved in apoptotic cell death. There are, however, a few areas that will need further studies in different research set up using both *in vitro* and *in vivo* tools: 1) is Pi induced apoptosis intrinsic or extrinsic or both? 2) how to link Pi caused ER stress to apoptosis? The results of our cell-based studies provide the template to establish further *in vitro* and *in vivo* studies to deepen our understanding of the underlying molecular mechanisms of high-phosphate-induced disease pathologies. Also, our results provide molecular signatures that have the potential to be either a biomarker and/or a therapeutic target to minimize high phosphate-induced cellular and tissue damages in diseases affecting various systems, such as cardiovascular to renal systems.

**In summary**, extracellular Pi rewired interwoven cell signaling networks bring about a broad spectrum of cytotoxicity, including aberrant proliferation, ER stress, EMT and cell death (Fig. 6A). Moderately high Pi promotes cell proliferation by synergistic activation of ERK and AKT pathways. At higher concentrations, Pi predominantly elicits cell death through MAPK and ER stress-mediated apoptosis, and EMT activated by TGF-β signaling (Fig. 6B). Our results form the basis of further studies on other cell-based and animal models to deepen our understanding of phosphate induced cellular responses, and to provide therapeutic clues to minimize phosphate toxicity-associated clinical complications.

## Materials and Methods

### Experimental Design

This study was designed to show Pi induced cellular responses so as to make insights into molecular mechanisms of phosphate toxicity in the cell model. We comprehensively investigate Pi related cellular events by exposing cells with increased amount of Pi for different durations. The XTT assay was used to study Pi effects on cell proliferation, survival and death. IF microscopy and WB were used to investigate Pi perturbed cell signaling. Inhibitors of AKT, MEK1/2, ER stress and JNK were used to study their inter-dependency as contributors to phosphotoxicity. All experiments were biologically repeated at least two times.

### Materials

Most of the reagents were purchased from Sigma-Aldrich (St. Louis, MO). HeLa cell line was a kind gift from Dr. Don Newmeyer (La Jolla Institute for Immunology, San Diego, CA). HEK293 cells, SH-SY5Y cells, Dimethylsulfoxide (DMSO), XTT Cell Proliferation Assay Kit and Universal Mycoplasma Detection Kit were purchased from ATCC (Manassas, VA). Dulbecco’s modified Eagle’s medium (DMEM), Dulbecco’s phosphate-buffered saline (DPBS), Ultrapure water, Nunc™ Lab-Tek™ II Chamber Slide™, EDTA (0.5 M), Halt™ Protease and Phosphatase Inhibitor Cocktail, UltraPure™ SDS Solution (10%), Pierce™ Rapid Gold BCA Protein Assay Kit, NuPAGE™ Bis-Tris 4-12% precast gels, NuPAGE™ MOPS SDS Running Buffer (20X), NuPAGE™ MES SDS Running Buffer (20X), Restore™ PLUS Western Blot Stripping Buffer, PageRuler™ Plus Prestained 10-250kDa Protein Ladder, ProLong™ Gold Antifade Mountant with DAPI, 16% Paraformaldehyde Aqueous Solution, Instant Nonfat Dry Milk, Trypsin-EDTA (0.25%), Fetal bovine serum (FBS) and L-Glutamine (200 mM) were obtained from Thermo Fisher Scientific (Carlsbad, CA). Nitrocellulose Membrane, Precision Plus Protein™ Dual Color Standards and Trans-Blot® Turbo™ RTA Midi Nitrocellulose Transfer Kit were purchased from Bio-Rad (Hercules, CA). Antibodies against Phospho-c-Raf (Ser338), Phospho-c-Raf (Ser259), c-Raf, Phospho-Akt (Ser473), Phospho-Akt (Thr308), Akt1, GAPDH, Phospho-p44/42 MAPK (Erk1/2) (Thr202/Tyr204), p44/42 MAPK (Erk1/2), Phospho-MEK1/2 (Ser217/221), MEK1/2, Phospho-SAPK/JNK (Thr183/Tyr185), SAPK/JNK, Phospho-p38 MAPK (Thr180/Tyr182), LC3A/B, BiP, Phospho-SMAD2 (Ser465/Ser467), Smad2, Phospho-RIP (Ser166), RIP, Phospho-MLKL (Ser358), MLKL, Cleaved Caspase-3 (Asp175), Caspase-3, Cleaved Caspase-8 (Asp384), Caspase-8, E-Cadherin, Vimentin, Snail, and chemicals including TPA, Staurosporine, Thapsigargin, Torin 1 were obtained from Cell Signaling Technology (Danvers, MA). Direct-Blot™ HRP anti-GAPDH and Direct-Blot™ HRP anti-β-actin, HRP Goat anti-mouse IgG, HRP donkey anti-rabbit IgG, HRP mouse anti-rat IgG antibodies, Western-Ready™ ECL Substrate Kit, and antibodies against ATF4, XBP-1s, p38 MAPK were purchased from BioLegend (San Diego, CA). NewBlot Stripping Buffer was purchased from LI-COR (Lincoln, NE). MK-2206, U-0126 and SP 600125 were obtained from Cayman Chemical (Ann Arbor, MI). Phalloidin-iFluor 488 Reagent was purchased from Abcam (Cambridge, MA).

### Cell culture and treatment

Human embryonic kidney cells (HEK293) and cervical cancer cells (HeLa) were maintained in DMEM (with 4.5g/L glucose, 1mM Pi) supplemented with 10% FBS. All cells were maintained 37 °C and 5% CO2. The cultured cells were routinely (every 10 passages of cells) quarantined for mycoplasma contamination using PCR-based mycoplasma detection kit.

Monosodium phosphate (NaH_2_PO_4_) was used as the source of Pi and sodium sulfate (Na_2_SO_4_) was used as the negative control. Cells were grown in regular DMEM containing 1mM Pi and 10% FBS and treated with an increase amount of Pi for 8hrs, 24hrs or 48hrs. For the inhibitors (MK-2206, 4-PBA, U-0126 and SP 600125) treatment, the cells were pre-treated with those compounds at indicated concentrations for 1hr followed by Pi treatment at different concentrations in the presence of inhibitors for 24hrs. Each treatment was repeated at least twice.

### XTT assay

The XTT assay was performed according to manufacturer’s instructions. Briefly, the optimal numbers (0.5-1 x10^4^) of cells were seeded into 96 well plates in triplicate. Cells were grown in complete medium for 24 hrs. After exposed to increased amounts of Pi for 24hrs, activated-XTT solution was added, and the plate will be returned to CO_2_ incubator for 3hrs. The absorbance was measured using Epoch 2 Microplate Spectrophotometer (BioTek Instruments, Inc., Winooski, VT). The average of specific absorbance from at least 3 biological replicates was normalized to 1mM Pi treated group. The relative absorbance value (fold change) was plotted as the mean absorbance ± SEM against the concentration of Pi or sulfate using Microsoft Excel 2010 software (Redmond, WA).

### IF Microscopy

HEK293 cells were grown in 8-well Chamber Slide to 80% confluence, and were exposed to increased Pi for 24hrs. Then, the cells were fixed in 4% paraformaldehyde in PBS for 15min followed by permeabilization with 0.5% Triton x-100 in PBS for 5min. Actin filaments were labeled with 1:2000 diluted Phalloidin-iFluor 488 reagents for 1h. Nuclei were counterstained with DAPI to analyze the apoptotic morphology. The image was captured with a 40X objective using Olympus IX53 fluorescence microscope (Shinjuku, Tokyo, Japan).

### Cell protein preparation

The cells were scraped in total lysis buffer (50 mM Tris-HCl, pH 7.4, 150 mM NaCl, 0.25% Sodium deoxycholate, 1.0% NP-40, 0.1% SDS) supplemented with 1X Protease and Phosphatase Inhibitor Cocktail. Protein concentration was measured by Rapid Gold BCA Protein Assay Kit according to manufacturer’s protocol.

### Electrophoresis and WB

A total of 15-30ug extracted protein was resolved by 4-12% Bis-Tris gel electrophoresis and transferred to nitrocellulose membrane. After blocking with 5% non-fat milk, the membrane was probed with diluted primary antibodies used at the manufacturer’s recommended concentrations. Proteins were visualized using HRP conjugated secondary antibodies and chemiluminescence detection. The blot was stripped and re-probed using NewBlot Stripping Buffer (LI-COR) or Restore™ PLUS Western Blot Stripping Buffer (Thermo Fisher). The images were captured by LI-COR C-Digit or Odyssey imaging systems (Lincoln, NE). The Ponceau S Stain of membrane or housekeeping genes (such as GAPDH and actin) expression level was used as a loading control. Blot quantification was done by densitometry image analysis using Image Studio Lite (V5.2) and Empiria Studio (V1.1) software (LI-COR, Lincoln, NE).

### Statistical Analysis

Data are expressed as means± SEM. Unpaired *Student t*-test was used to compare means between control and high Pi treated groups. A paired *t*-test was used to compare means between non-treated and inhibitors treated groups at indicated concentrations of Pi. A value of *P* < 0.05 was considered statistically significant. GraphPad Prism 8.0 software programs were used to plot the data and perform the statistics.

## Acknowledgments

We are grateful to Dr. Yanfei Qi, Dr. Alice Hudder, Dr. Diana Speelman and Dr. Randy Kulesza (Lake Erie College of Osteopathic Medicine, Erie, PA) for their willingness to share their lab space, instruments and biological reagents with our group. We also appreciate Rebecca Hetz’s assistance in Western blot experiments and cell culture.

## Funding

This work was supported by LECOM internal Seed Grants (funding opportunity number: COM-18-19) and Research foundation.

## Author contribution

P.H. led the whole study including study design, performing experiments and data analysis, results interpretation, and manuscript writing. O.M.C. assisted in the XTT assays, Western blot experiments and critically reviewed the manuscript. J.F. assisted in Western blot experiments and data analysis. N.E. assisted in Western blot experiments and critically reviewed the manuscript. M.S.R. contributed to the study design, data discussion and editing of the manuscript.

## Competing interests

The authors declare that they have no competing interests.

## Data and materials availability

All data required to assess the conclusions in the manuscript are present in the manuscript and/or the Supplementary Materials. Additional data related to this paper may be requested from P.H.

## Supplementary Materials

**Fig. S1.**
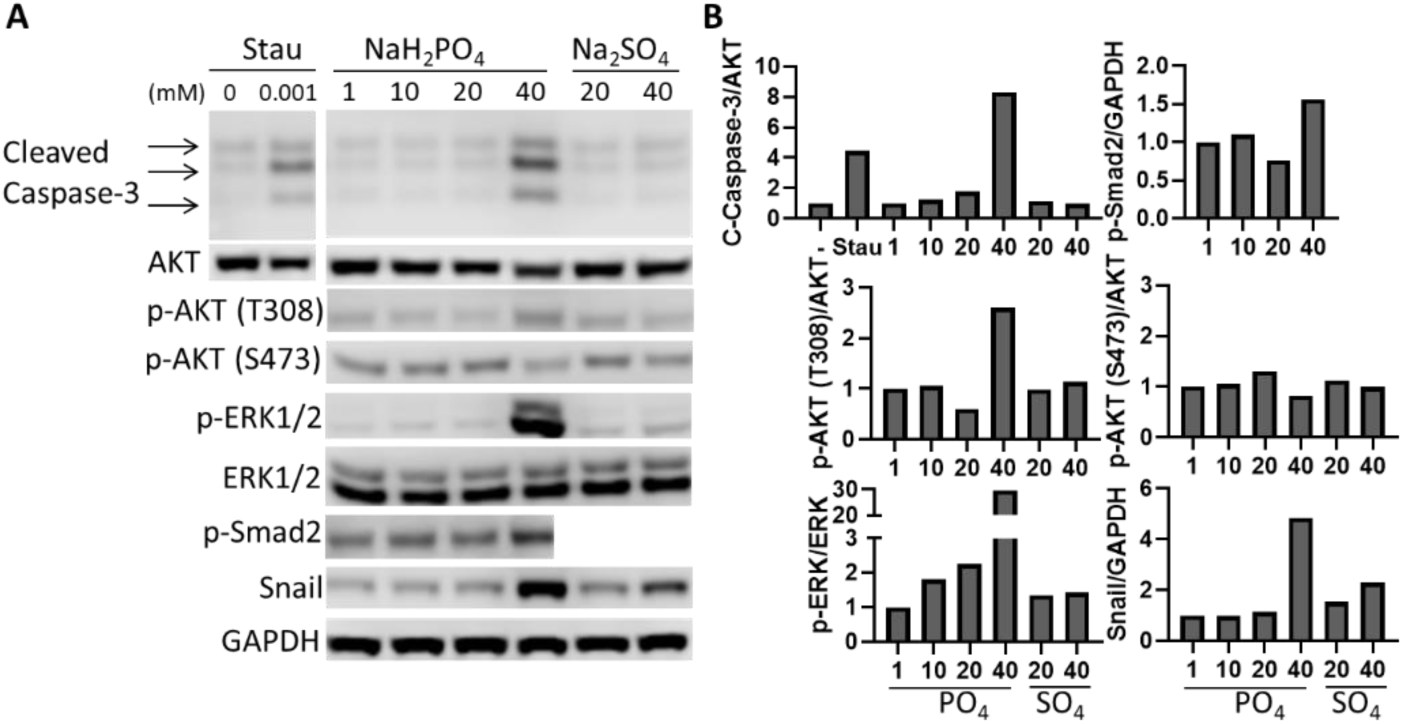
High concentration of Pi triggers apoptosis and EMT as early as 8 hrs in HEK293 cells. WB analysis **(A)** shows 40mM NaH_2_PO_4_ (Pi), compared to Na2SO_4_ (negative control), could promote apoptosis and EMT after 8hrs treatment. The high concentration of Pi also activated ERK1/2 and AKT(T308) phosphorylation. GAPDH was used as loading control. Bands quantification **(B)** was done by densitometry image analysis using Image Studio Lite software. The ratios of phosphorylated /Pan for each target or target protein/GAPDH is divided by that of the control (1mM Pi) to obtain the relative value for each treatment.

**Fig. S2.**
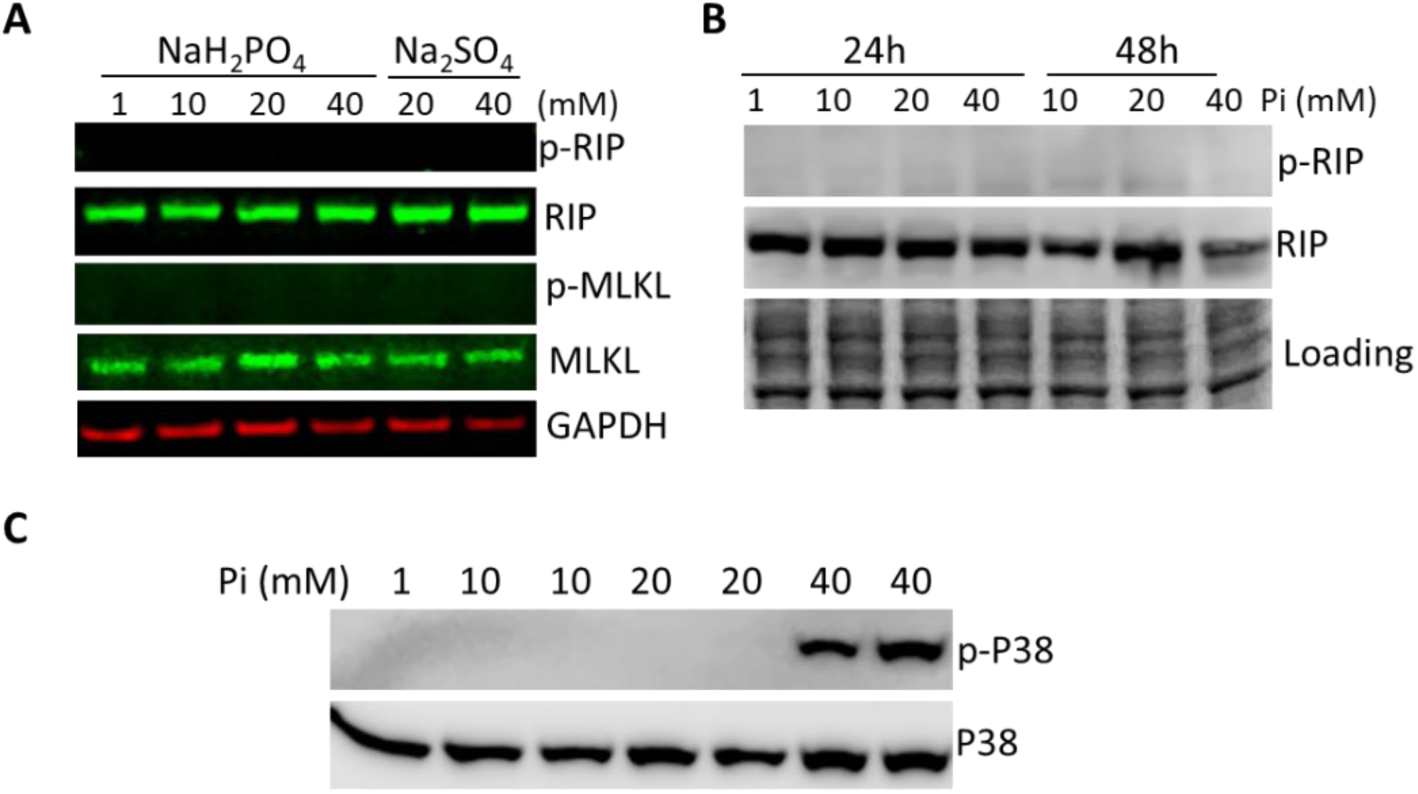
High concentration of Pi does not cause necroptosis (A and B), but activates p38 (C) in HEK293 cells.

**Fig. S3.**
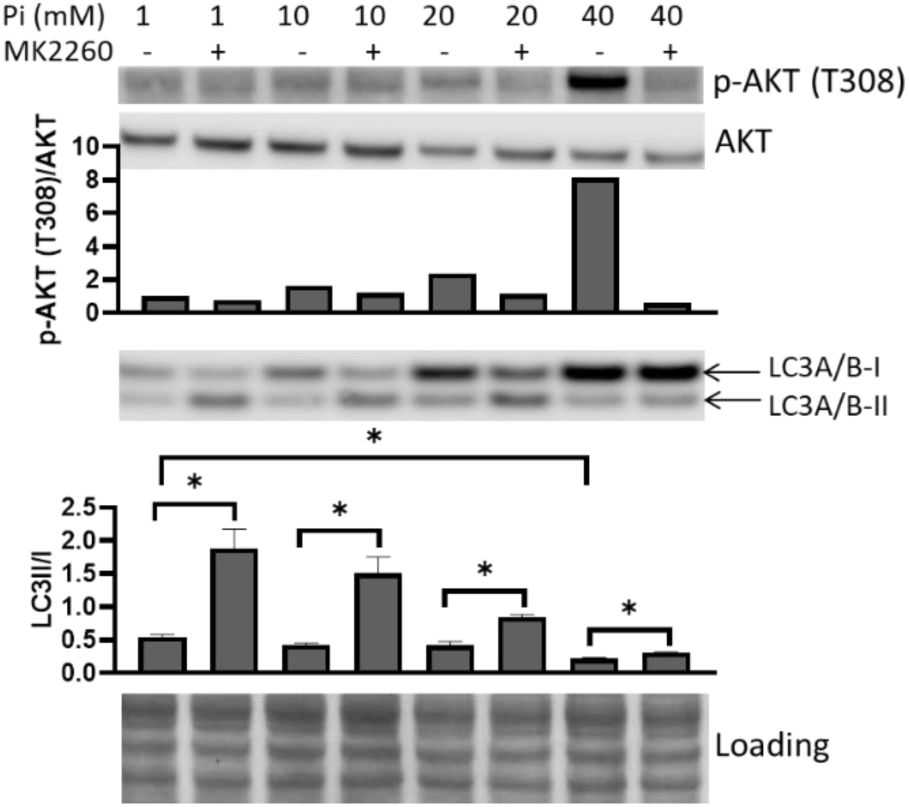
MK-2206 unleashes high Pi inhibited autophagy in HeLa cells. WB analysis shows MK-2206 alleviated Pi suppressed autophagy (reduced ratio of LC-II/LC-I). Data with error bars represent means ± SEM from at least three independent experiments. Unpaired *Student t*-test was used to compare means between 1mM and higher concentrations of Pi treated groups. Paired *t*-test was used to compare means between non-treated and MK-2206 treated groups at indicated concentrations of Pi. \**P* < 0.05.

**Fig. S4.**
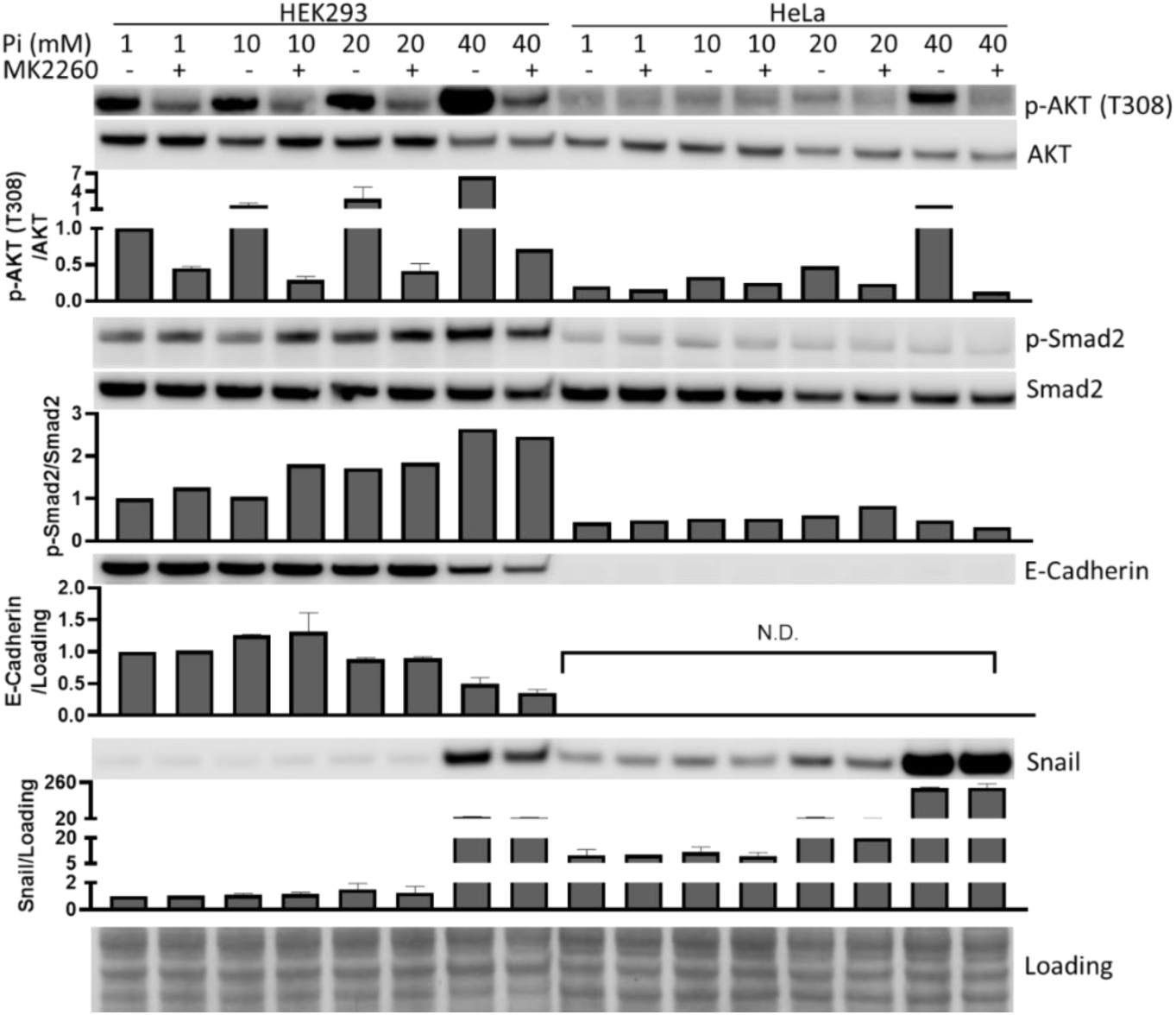
MK-2206 does not affect EMT in HEK293 and HeLa cells. WB analysis shows MK-2206 does not significantly change EMT markers’ level. Data with error bars represent means ± SEM from at least three independent experiments. Paired *t*-test was used to compare means between non-treated and MK-2206 treated groups at indicated concentrations of Pi. N.D., not determined.

**Fig. S5.**
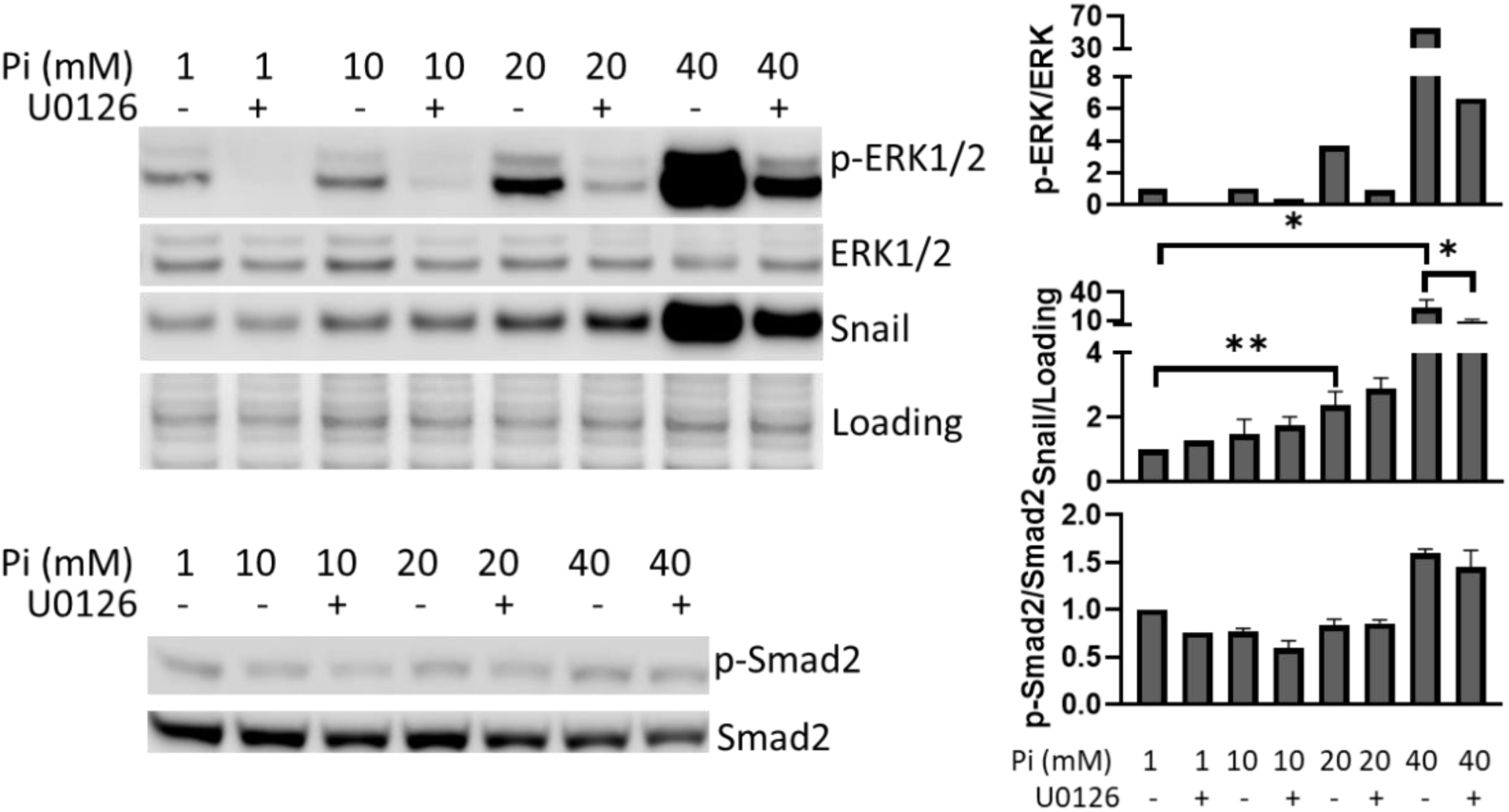
U-0126 represses Pi stimulated Snail up-regulation in HeLa cells. WB analysis shows U-0126 partially prevented 40mM Pi induced Snail up-regulation but had no effect on phosphorylation of Smad2. Data with error bars represent means ± SEM from at least three independent experiments. Unpaired *Student t*-test was used to compare means between 1mM and higher concentrations of Pi treated groups. Paired *t*-test was used to compare means between non-treated and U-0126 treated groups at indicated concentrations of Pi. \**P* < 0.05.

## References

Beck GR (2003) Inorganic phosphate as a signaling molecule in osteoblast differentiation. J Cell Biochem 90: 234–43

Beck GR, Jr., Knecht N (2003) Osteopontin regulation by inorganic phosphate is ERK1/2-, protein kinase C-, and proteasome-dependent. J Biol Chem 278: 41921–9

Brown RB, Razzaque MS (2018) Phosphate toxicity and tumorigenesis. Biochim Biophys Acta Rev Cancer 1869: 303–309

Cagnol S, Chambard JC (2010) ERK and cell death: mechanisms of ERK-induced cell death--apoptosis, autophagy and senescence. FEBS J 277: 2–21

Camalier CE, Yi M, Yu LR, Hood BL, Conrads KA, Lee YJ, Lin Y, Garneys LM, Bouloux GF, Young MR, Veenstra TD, Stephens RM, Colburn NH, Conrads TP, Beck GR (2013) An integrated understanding of the physiological response to elevated extracellular phosphate. J Cell Physiol 228: 1536–50

Camalier CE, Young MR, Bobe G, Perella CM, Colburn NH, Beck GR (2010) Elevated phosphate activates N-ras and promotes cell transformation and skin tumorigenesis. Cancer Prev Res (Phila) 3: 359–70

Cancela AL, Santos RD, Titan SM, Goldenstein PT, Rochitte CE, Lemos PA, dos Reis LM, Graciolli FG, Jorgetti V, Moyses RM (2012) Phosphorus is associated with coronary artery disease in patients with preserved renal function. PLoS One 7: e36883

Chang SH, Yu KN, Lee YS, An GH, Beck GR, Colburn NH, Lee KH, Cho MH (2006) Elevated inorganic phosphate stimulates Akt-ERK1/2-Mnk1 signaling in human lung cells. Am J Respir Cell Mol Biol 35: 528–39

Conrads KA, Yi M, Simpson KA, Lucas DA, Camalier CE, Yu LR, Veenstra TD, Stephens RM, Conrads TP, Beck GR (2005) A combined proteome and microarray investigation of inorganic phosphate-induced pre-osteoblast cells. Mol Cell Proteomics 4: 1284–96

Di Marco GS, Hausberg M, Hillebrand U, Rustemeyer P, Wittkowski W, Lang D, Pavenstädt H (2008) Increased inorganic phosphate induces human endothelial cell apoptosis in vitro. Am J Physiol Renal Physiol 294: F1381–7

Dibble CC, Cantley LC (2015) Regulation of mTORC1 by PI3K signaling. Trends Cell Biol 25: 545–55

Doi S, Zou Y, Togao O, Pastor JV, John GB, Wang L, Shiizaki K, Gotschall R, Schiavi S, Yorioka N, Takahashi M, Boothman DA, Kuro-o M (2011) Klotho inhibits transforming growth factor-beta1 (TGF-beta1) signaling and suppresses renal fibrosis and cancer metastasis in mice. J Biol Chem 286: 8655–65

Favata MF, Horiuchi KY, Manos EJ, Daulerio AJ, Stradley DA, Feeser WS, Van Dyk DE, Pitts WJ, Earl RA, Hobbs F, Copeland RA, Magolda RL, Scherle PA, Trzaskos JM (1998) Identification of a novel inhibitor of mitogen-activated protein kinase kinase. J Biol Chem 273: 18623–32

Fruman DA, Chiu H, Hopkins BD, Bagrodia S, Cantley LC, Abraham RT (2017) The PI3K Pathway in Human Disease. Cell 170: 605–635

Giachelli CM (2009) The emerging role of phosphate in vascular calcification. Kidney Int 75: 890–7

Goodson JM, Shi P, Razzaque MS (2019) Dietary phosphorus enhances inflammatory response: A study of human gingivitis. J Steroid Biochem Mol Biol

Gui T, Sun Y, Shimokado A, Muragaki Y (2012) The Roles of Mitogen-Activated Protein Kinase Pathways in TGF-beta-Induced Epithelial-Mesenchymal Transition. J Signal Transduct 2012: 289243

Gunther C, Martini E, Wittkopf N, Amann K, Weigmann B, Neumann H, Waldner MJ, Hedrick SM, Tenzer S, Neurath MF, Becker C (2011) Caspase-8 regulates TNF-alpha-induced epithelial necroptosis and terminal ileitis. Nature 477: 335–9

Hanahan D, Weinberg RA (2011) Hallmarks of cancer: the next generation. Cell 144: 646–74

Hetz C, Martinon F, Rodriguez D, Glimcher LH (2011) The unfolded protein response: integrating stress signals through the stress sensor IRE1alpha. Physiol Rev 91: 1219–43

Hirai H, Sootome H, Nakatsuru Y, Miyama K, Taguchi S, Tsujioka K, Ueno Y, Hatch H, Majumder PK, Pan BS, Kotani H (2010) MK-2206, an allosteric Akt inhibitor, enhances antitumor efficacy by standard chemotherapeutic agents or molecular targeted drugs in vitro and in vivo. Mol Cancer Ther 9: 1956–67

Iurlaro R, Munoz-Pinedo C (2016) Cell death induced by endoplasmic reticulum stress. FEBS J 283: 2640–52

Jiang Z, Fan Q, Zhang Z, Zou Y, Cai R, Wang Q, Zuo Y, Cheng J (2012) SENP1 deficiency promotes ER stress-induced apoptosis by increasing XBP1 SUMOylation. Cell Cycle 11: 1118–22

Jin H, Xu CX, Lim HT, Park SJ, Shin JY, Chung YS, Park SC, Chang SH, Youn HJ, Lee KH, Lee YS, Ha YC, Chae CH, Beck GR, Cho MH (2009) High dietary inorganic phosphate increases lung tumorigenesis and alters Akt signaling. Am J Respir Crit Care Med 179: 59–68

Julien M, Magne D, Masson M, Rolli-Derkinderen M, Chassande O, Cario-Toumaniantz C, Cherel Y, Weiss P, Guicheux J (2007) Phosphate stimulates matrix Gla protein expression in chondrocytes through the extracellular signal regulated kinase signaling pathway. Endocrinology 148: 530–7

Kalantar-Zadeh K, Gutekunst L, Mehrotra R, Kovesdy CP, Bross R, Shinaberger CS, Noori N, Hirschberg R, Benner D, Nissenson AR, Kopple JD (2010) Understanding sources of dietary phosphorus in the treatment of patients with chronic kidney disease. Clin J Am Soc Nephrol 5: 519–30

Kanatani M, Sugimoto T, Kano J, Kanzawa M, Chihara K (2003) Effect of high phosphate concentration on osteoclast differentiation as well as bone-resorbing activity. J Cell Physiol 196: 180–9

Kimata M, Michigami T, Tachikawa K, Okada T, Koshimizu T, Yamazaki M, Kogo M, Ozono K (2010) Signaling of extracellular inorganic phosphate up-regulates cyclin D1 expression in proliferating chondrocytes via the Na+/Pi cotransporter Pit-1 and Raf/MEK/ERK pathway. Bone 47: 938–47

Klionsky DJ, Abdelmohsen K, Abe A, Abedin MJ, Abeliovich H, Acevedo Arozena A, Adachi H, Adams CM, Adams PD, Adeli K, Adhihetty PJ, Adler SG, Agam G, Agarwal R, Aghi MK, Agnello M, Agostinis P, Aguilar PV, Aguirre-Ghiso J, Airoldi EM et al. (2016) Guidelines for the use and interpretation of assays for monitoring autophagy (3rd edition). Autophagy 12: 1–222

Kuro OM (2011) Phosphate and Klotho. Kidney Int Suppl: S20–3

Lavoie H, Therrien M (2015) Regulation of RAF protein kinases in ERK signalling. Nature reviews Molecular cell biology 16: 281–98

Lee S, Kim JE, Hong SH, Lee AY, Park EJ, Seo HW, Chae C, Doble P, Bishop D, Cho MH (2015) High Inorganic Phosphate Intake Promotes Tumorigenesis at Early Stages in a Mouse Model of Lung Cancer. PLoS One 10: e0135582

Lin Y, McKinnon KE, Ha SW, Beck GR (2015) Inorganic phosphate induces cancer cell mediated angiogenesis dependent on forkhead box protein C2 (FOXC2) regulated osteopontin expression. Mol Carcinog 54: 926–34

Mancini FR, Affret A, Dow C, Balkau B, Clavel-Chapelon F, Bonnet F, Boutron-Ruault MC, Fagherazzi G (2018) High dietary phosphorus intake is associated with an increased risk of type 2 diabetes in the large prospective E3N cohort study. Clin Nutr 37: 1625–1630

Manning BD, Toker A (2017) AKT/PKB Signaling: Navigating the Network. Cell 169: 381–405

Marks J, Debnam ES, Unwin RJ (2013) The role of the gastrointestinal tract in phosphate homeostasis in health and chronic kidney disease. Curr Opin Nephrol Hypertens 22: 481–7

Michigami T, Kawai M, Yamazaki M, Ozono K (2018) Phosphate as a Signaling Molecule and Its Sensing Mechanism. Physiol Rev 98: 2317–2348

Morrison DK (2012) MAP kinase pathways. Cold Spring Harb Perspect Biol 4

Nakatani T, Sarraj B, Ohnishi M, Densmore MJ, Taguchi T, Goetz R, Mohammadi M, Lanske B, Razzaque MS (2009) In vivo genetic evidence for klotho-dependent, fibroblast growth factor 23 (Fgf23) -mediated regulation of systemic phosphate homeostasis. FASEB J 23: 433–41

Nishino J, Yamazaki M, Kawai M, Tachikawa K, Yamamoto K, Miyagawa K, Kogo M, Ozono K, Michigami T (2017) Extracellular Phosphate Induces the Expression of Dentin Matrix Protein 1 Through the FGF Receptor in Osteoblasts. J Cell Biochem 118: 1151–1163

Qi X, Hosoi T, Okuma Y, Kaneko M, Nomura Y (2004) Sodium 4-phenylbutyrate protects against cerebral ischemic injury. Mol Pharmacol 66: 899–908

Roussanne MC, Lieberherr M, Souberbielle JC, Sarfati E, Drüeke T, Bourdeau A (2001) Human parathyroid cell proliferation in response to calcium, NPS R-467, calcitriol and phosphate. Eur J Clin Invest 31: 610–6

Saxton RA, Sabatini DM (2017) mTOR Signaling in Growth, Metabolism, and Disease. Cell 168: 960–976

Thoreen CC, Kang SA, Chang JW, Liu Q, Zhang J, Gao Y, Reichling LJ, Sim T, Sabatini DM, Gray NS (2009) An ATP-competitive mammalian target of rapamycin inhibitor reveals rapamycin-resistant functions of mTORC1. J Biol Chem 284: 8023–32

Tonelli M, Sacks F, Pfeffer M, Gao Z, Curhan G, Cholesterol, Recurrent Events Trial I (2005) Relation between serum phosphate level and cardiovascular event rate in people with coronary disease. Circulation 112: 2627–33

Walter P, Ron D (2011) The unfolded protein response: from stress pathway to homeostatic regulation. Science 334: 1081–6

Wolf I, Levanon-Cohen S, Bose S, Ligumsky H, Sredni B, Kanety H, Kuro-o M, Karlan B, Kaufman B, Koeffler HP, Rubinek T (2008) Klotho: a tumor suppressor and a modulator of the IGF-1 and FGF pathways in human breast cancer. Oncogene 27: 7094–105

Yamazaki M, Ozono K, Okada T, Tachikawa K, Kondou H, Ohata Y, Michigami T (2010) Both FGF23 and extracellular phosphate activate Raf/MEK/ERK pathway via FGF receptors in HEK293 cells. J Cell Biochem 111: 1210–21

Yoshikawa R, Yamamoto H, Nakahashi O, Kagawa T, Tajiri M, Nakao M, Fukuda S, Arai H, Masuda M, Iwano M, Takeda E, Taketani Y (2018) The age-related changes of dietary phosphate responsiveness in plasma 1,25-dihydroxyvitamin D levels and renal Cyp27b1 and Cyp24a1 gene expression is associated with renal α-Klotho gene expression in mice. J Clin Biochem Nutr 62: 68–74

Zhang W, Liu HT (2002) MAPK signal pathways in the regulation of cell proliferation in mammalian cells. Cell Res 12: 9–18

Zhang YY, Yang M, Bao JF, Gu LJ, Yu HL, Yuan WJ (2018) Phosphate stimulates myotube atrophy through autophagy activation: evidence of hyperphosphatemia contributing to skeletal muscle wasting in chronic kidney disease. BMC Nephrol 19: 45

